# DrugCombDB: a comprehensive database of drug combinations toward network medicine and combination therapy

**DOI:** 10.1101/477547

**Authors:** Lei Deng, Bo Zou, Wenhao Zhang, Hui Liu

**Affiliations:** School of Software, Central South University, Changsha, 410075 China; Lab of Information Management, Changzhou University, 213164 China

## Abstract

Drug combinations have demonstrated high efficacy and low adverse side effects compared to single drug administrations in cancer therapies, and thus draw intensive attentions from researchers and pharmaceutical enterprises. Thanks to the fast development of high-throughput screening (HTS) methods, the amount of available drug combination datasets has tremendously increased. However, existing drug combination databases are lack of indications of the drug combinations and quantitative dose-responses. Therefore, there is an urgent need for a comprehensive database that is crucial to both experimental and computational screening of drug combinations. In this paper, we present DrugCombDB, a comprehensive database dedicated to integrating drug combinations from various data sources. Concretely, the data sources include 1) high-throughput screening assays of drug combinations, 2) external databases, and 3) manual curations from PubMed literature. In total, DrugCombDB includes 1,127,969 experimental data points with quantitative dose response and concentrations of drug combinations covering 561 unique drugs and 104 human cancer cell lines, and 1,875 FDA approved or literature-supported drug combinations. In particular, we adopted the zero interaction potency (ZIP) model [2] to compute the scores determining the synergy or antagonism of two drugs. To facilitate the downstream usage of our data resource, we prepared multiple datasets that are ready for building prediction models of classification and regression analysis. A website with user-friendly data visualization is provided to help users access the wealth of data. Users can input a drug of interest to retrieve associated drug combinations, together with the supporting evidence sources and drug targets. Our database is available at http://drugcombdb.denglab.org/.

## Background

Although targeted drugs have led to remarkable advances in the treatment of cancer patients, their clinical benefits to tumor therapies are greatly limited due to intrinsic and acquired resistance of cancer cells against such drugs. The root cause of development to drug resistance lie in that compensatory kinases and pathways become activated and maintain the growth and survival of tumor cells. Drug combinations have demonstrated great advantage to improve efficacy and overcome resistance for treating complex and refractory diseases, compared to single drug administrations in cancer therapies, and thus draw increasing attentions from researchers and pharmaceutical enterprises. Despite the increasing successes of combination drugs in inhibiting cancer cell proliferation, most of them are discovered by clinical experience or by occasional chances. So, there is an urgent demand for rational and systematic methodology to screen cancer-specific and sensitive combinatorial drugs for cancer therapy. With insight gained by the understanding of pathway interdependencies that are critical for cancer cell proliferation and survival in a specific cancer type, researchers are able to design multiple agents to synergistically inhibit signaling pathways. However, the wet-lab experiments currently used to dissect the cellular mechanism of cascade signal transduction and signaling network.

Thanks to the fast development of high-throughput screening (HTS) methods, it is possible to systematically evaluate the pairwise combinations from a large number of both approved and investigational chemical compounds. As a result, the amount of available drug combination datasets has tremendously increased, which can benefit the researchers a lot. However, existing database DCDB [1], which has not been updated since 2014, covers only 1,363 drug combination annotations extracted from FDA orange books and records of clinical trials using the text-mining technique, which are lack of indications of the drug combinations and quantitative dose-responses. Therefore, there is an urgent need for a comprehensive database that is crucial to both experimental and computational screening of drug combinations.

In this paper, we present DrugCombDB, a comprehensive database dedicated to integrating drug combinations from various data sources. Concretely, the data sources include 1) high-throughput screening assays of drug combinations, 2) external databases, and 3) manual curations from PubMed literature. In total, DrugCombDB includes 1,127,969 experimental data points with quantitative dose response and concentrations of drug combinations covering 561 unique drugs and 104 human cancer cell lines, and 1,875 FDA approved or literature-supported drug combinations. In particular, we adopted the zero interaction potency (ZIP) model [2] to compute the scores determining the synergy or antagonism of two drugs. To facilitate the downstream usage of our data resource, we prepared multiple datasets that are ready for building prediction models of classification and regression analysis. Moreover, a website with user-friendly data visualization is developed to help users access the wealth of data. Users can input a drug of interest to retrieve associated drug combinations, together with the supporting evidence sources and drug targets. The dose responses and drugs concentrations with respect to cancer cell lines are displayed in interactive scatter plots.

To the best of our knowledge, DrugCombDB is the first comprehensive database with the largest number of drug combinations to date. We believe it would greatly facilitate and accelerate the discovery of novel synergistic drugs for the therapy of complex diseases, especially for the cancers developed drug resistance.

## Data resources

### HTS assays

The main data source of DrugCombDB comes from high-throughput screening assays that are released by publications concentrated on combinational therapies. We have conducted careful publication retrieval to collect experimental data sets via PubMed and search engine. As a result, three large-scale wet-lab experiments carried to explore efficacy of drug combination are collected from related publications [3,4]. In these experiments, high-throughput screening assays are applied to identify the combinatorial efficacy (synergy, additivity and antagonism) between different drugs, in which the quantitative dose responses of cancer cells to different drug combinations and different concentrations are recorded.

Mohammad et al. aimed to explore adaptive resistance of melanoma cells to RAF inhibition [3]. By performing dye-based imaging assays, the dose responses of BRAFV600E melanoma cells in response to Vemurafenib with another compound were monitored. Viability and apoptosis were scored to measure the resulting states of BRAF melanoma cells. We collected 620 experimental data points in total, in which Vemurafenib was in combination with 37 different compounds at different concentrations.

Similarly, a high throughput screening platform is adopted to discover therapeutic combinations for the activated B-celllike subtype (ABC) of diffuse large B cell lymphoma (DLBCL) [5]. The therapeutic efficacy between different drugs in combination with Brutons tyrosine kinase inhibitor ibrutinib are measured. Ibrutinib is designed to target the chronic active B-cell receptor signaling pathway that characterizes ABC DLBCL. In total, 466 different agents were evaluated in combination with ibrutinib using 6 × 6 dose-response blocks. As a result, 16,776 experimental data points were integrated into DrugCombDB.

Jennifer et al. presented an unbiased oncology compound screening to identify novel combination strategies [4]. The high-throughput screening for combinatorial drugs was performed on the fully automated GNF PolyTarget robotic platform, where cells were treated with a 4 by 4 matrix of drug concentrations. Based on the cell viability measured by using CellTiter-Glo cell viability reagent (Promega), the highest single agent (HSA) and Bliss independence models were applied to determine the combinatorial efficacy. Totally, this assay yield to 22,737 data points of 583 pairwise drug combinations over 39 diverse cancer cell lines.

Moreover, a large amount of organized experimental points from national institutes of health (NIH) database supported by national cancer institute (NCI), which is also devoted to helping researchers from related fields to do analytical work, were integrated into DrugCombDB. Thanks for the persevering effort of NCI, we are able to download the data set that is available on NCI website [7]. This assay was conducted within 3×3 dose-response matrix containing 82,209 pairs of drug combinations. Consequently, 739,881 data points were integrated to DrugCombDB.

In summary, DrugCombDB contains 1,126,109 experimental data points covering 105,449 pairwise drug combinations, 561 unique drugs and 104 cancer cell lines.

### Text Mining

To extend the coverage of DrugCombDB, text mining tools are employed to extract drug combinations from literature. Using drug combinations, combination drug, combinatorial drugs, and synergistic drugs as query keywords, we searched the PubMed database and obtained 922 distinct publications with titles including one of these keywords. These publications focus on the therapeutic effect of various drug combinations using in vitro models and flow cytometry. Subsequently, we adopted PubTator [5], a web-based tool facilitating manual literature curation through powerful text-mining techniques, to annotate the abstracts of the publications. Taking the PubMed ID list of the filtered publications as the input, PubTator can mark the discriminative concepts such as gene, chemical, disease, species and mutations in different colors. Consequently, we manually check the titles and abstracts with highlighted concepts, and identify drug combinations that have demonstrated therapeutic efficacy in certain cancer cells in these publications.

### Three-drug combinations

Due to the limitations of experimental capability, the high-throughput assays were generally conducted within dose-response matrixes, which can only accommodate two drugs. Accordingly, double-drug treatments are the research focus in drug screening. However, complicated drug combinations mean more targets, which can improve efficacy and overcome resistance for treating complex and refractory diseases. During the process of collecting data, especially in text-mining, we also collected many combinations of multiple drugs evaluated by biochemical experiments or clinical trials. In order to expand the scale of DrugCombDB, we also integrated these data into our database for enhanced functionalities.

## Data Normalization

As the drug combinations in DrugCombDB were collected from different types of sources, including biochemical assays, high-throughput screening experiments, other related databases and text mining results, the dose response values vary from different experimental protocol and platforms. To facilitate usage of our database, we normalized the dose response values to give coincident and comparable therapeutic efficacy over different data sets. Considering that the cell viability and the apoptosis rate upon treatments are the most used measures in drug combination assays, we introduced the normalized inhibition rate of cancer cell to drug treatments as the uniform measure, using min-max normalization that is defined below:

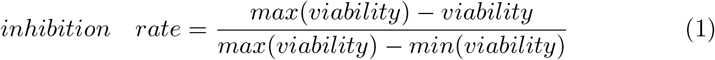

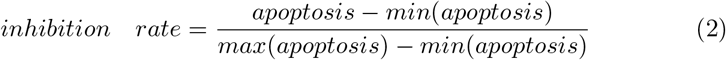

As a result, 1 represents the most synergistic efficacy and 0 represents the most antagonistic efficacy. The normalized inhibition rates range from 0 to 1, which is favorable for regression analysis. It is worth noting that, for each tested drug combination, HTS assays yield to a number of experimental data points comprised of a dose response matrix corresponding to different concentration combinations. Together with the drug concentrations, the normalized inhibition rates greatly expand the volume of training data for regression analysis.

## Classification of synergism and antagonism

It is well-known that the efficacy of drug combinations is classified as synergistic or antagonistic, depending on that cancer cells are inhibited from or accelerated to proliferation than the additivity efficacy of independent treatment of two drugs. The synergistic or antagonistic effect can be determined by the deviation of the dose response curves from the expected effect calculated based on a reference model of Loewe additivity or bliss independence. When the percentage of inhibited or killed cancer cells is greater than expected, then the drug combinations are classified as synergistic. On the other hand, antagonism is determined when the drug combination produces effect worse than expected. Although the normalized inhibition rate mentioned above quantify the response of cancer cells to certain concentrations of drug combinations, the cutoff of synergy and antagonism cannot be simply determined. To build the training set for classification model, we proceed to process these data sets using comprehensive model that takes the whole dose response matrix into account.

The existing scoring models to quantify the efficacy of drug combinations, for example, highest single agent model and bliss independence model, were proposed originally for low-throughput drug combination experiments. When tackling large-scale doseresponse experiments with various dose pairs, the model assumptions are not capable for the complex patterns of drug interactions. To overcome the limitations, we adopt a novel reference model named zero interaction potency (ZIP), to further process our data sets. ZIP model has been demonstrated to capture the drug interaction relationships by comparing the change in the potency of the doseresponse curves between individual drugs and their combinations [3].

ZIP is a response surface model that combines the advantages of the Loewe and the Bliss models, which proposed a delta score to characterize the synergy landscape over the full dose-response matrix. The ZIP model assumes that two non-interacting drugs are expected to incur minimal changes in their dosere-sponse curves. A delta score is computed to quantify the deviation from the expectation of ZIP for a given dose pair and utilized the average delta over a doseresponse matrix as a summary interaction score for a drug combination. As a result, ZIP model is perfectly compatible with high-throughput drug combination screening data. One advantage over traditional model is that ZIP model has definite threshold derived from mathematical reasoning to determine the classification of drug combinations (synergy or antagonism). Formally, the drug combinations with ZIP score greater than 0 are classified as synergistic ones, otherwise antagonistic ones. Therefore, we computed the ZIP score by running SynergyFinder [2], a web application for analyzing drug combination dose-response matrix, for each data sets, and classify each drug combinations to synergy or antagonism according to its ZIP score.

## FUNCTIONALITIES

A website with user-friendly data visualization is provided to help users access the wealth of data. We have developed a few functional modules to assist downstream users explore the value of our website. The browser module is the fundamental section, where all drug combinations we collected can be displayed with different entries. Meanwhile, to help users make sufficient usage, more detailed information of the associated drugs is also displayed, including molecular weight and smile string, which can be retrieved on Stitch. The browser page is shown in Figure 1.

**Fig. 1.**
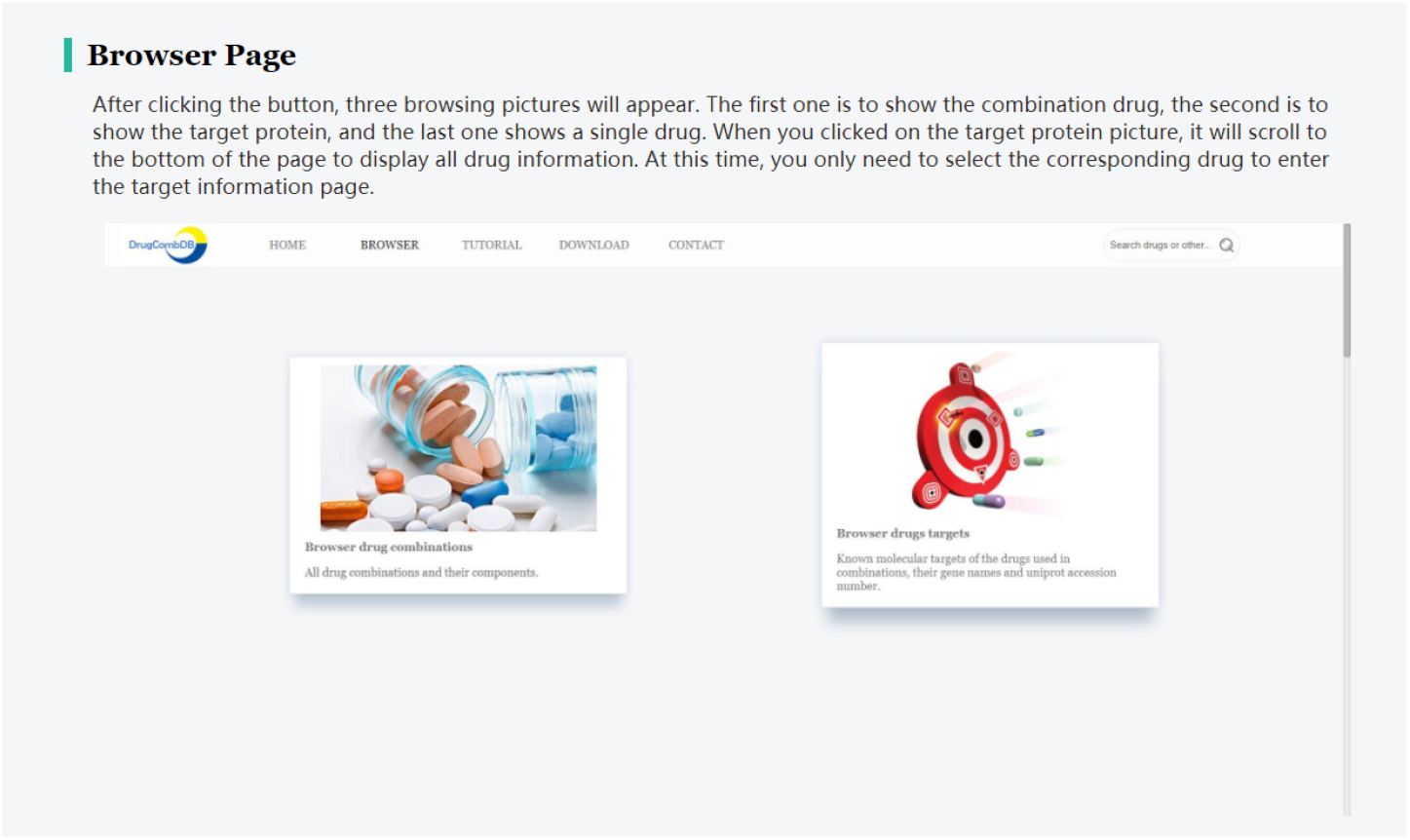
The predicted sensitivities of the combination drugs Lapatinib and Rapamycin.

The query module takes drug names as input to search for all related drug combinations containing the drug, and the query drug combinations with respect to cancer cell lines will be presented in the form of both interactive scatter plot and tabular viewers. In the scatter plot viewer, each dynamic point represents a pair of drug combination with concentrations, the horizontal axis corresponds to different cell lines targeted by the drug combination, and the the vertical axis indicates the normalized growth rate, as shown in Figure 2. In particular, the same drug combination has diverse concentrations, therefore, different kinds of drugs are marked with different colors for better distinction. Detailed information about the combinations can be displayed by clicking the hyperlinks of the scatters, where other original data about the combination assay and its source can be explored, as shown in Figure 3. In the tabular viewer, the participated drugs of the combination, corresponding concentrations, normalized dose response or referred to as growth, targeting cell line and its supporting literature source are displayed for each combination, as shown in Figure 4.

**Fig. 2.**
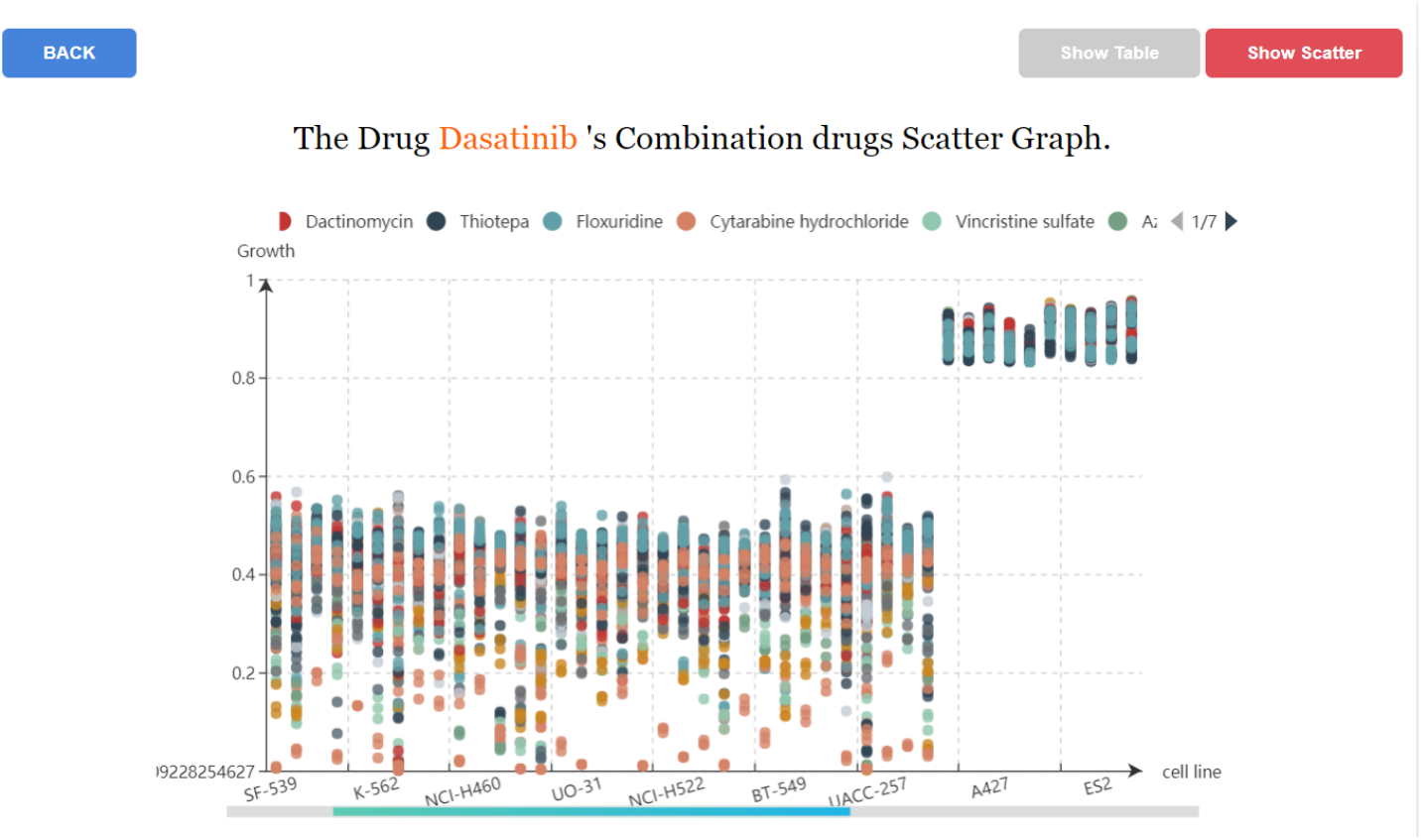
The predicted sensitivities of the combination drugs Lapatinib and Rapamycin.

**Fig. 3.**
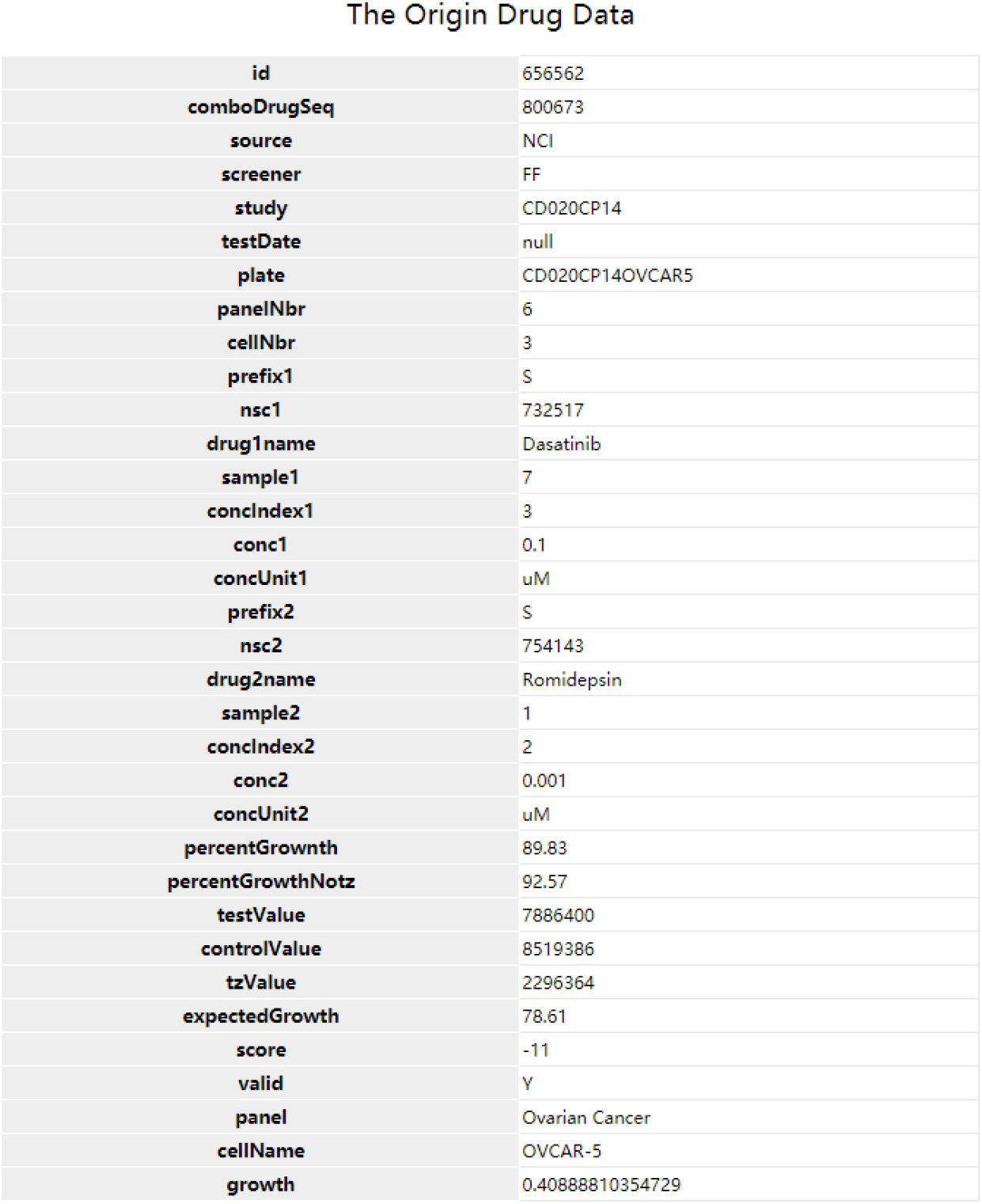
The predicted sensitivities of the combination drugs Lapatinib and Rapamycin.

**Fig. 4.**
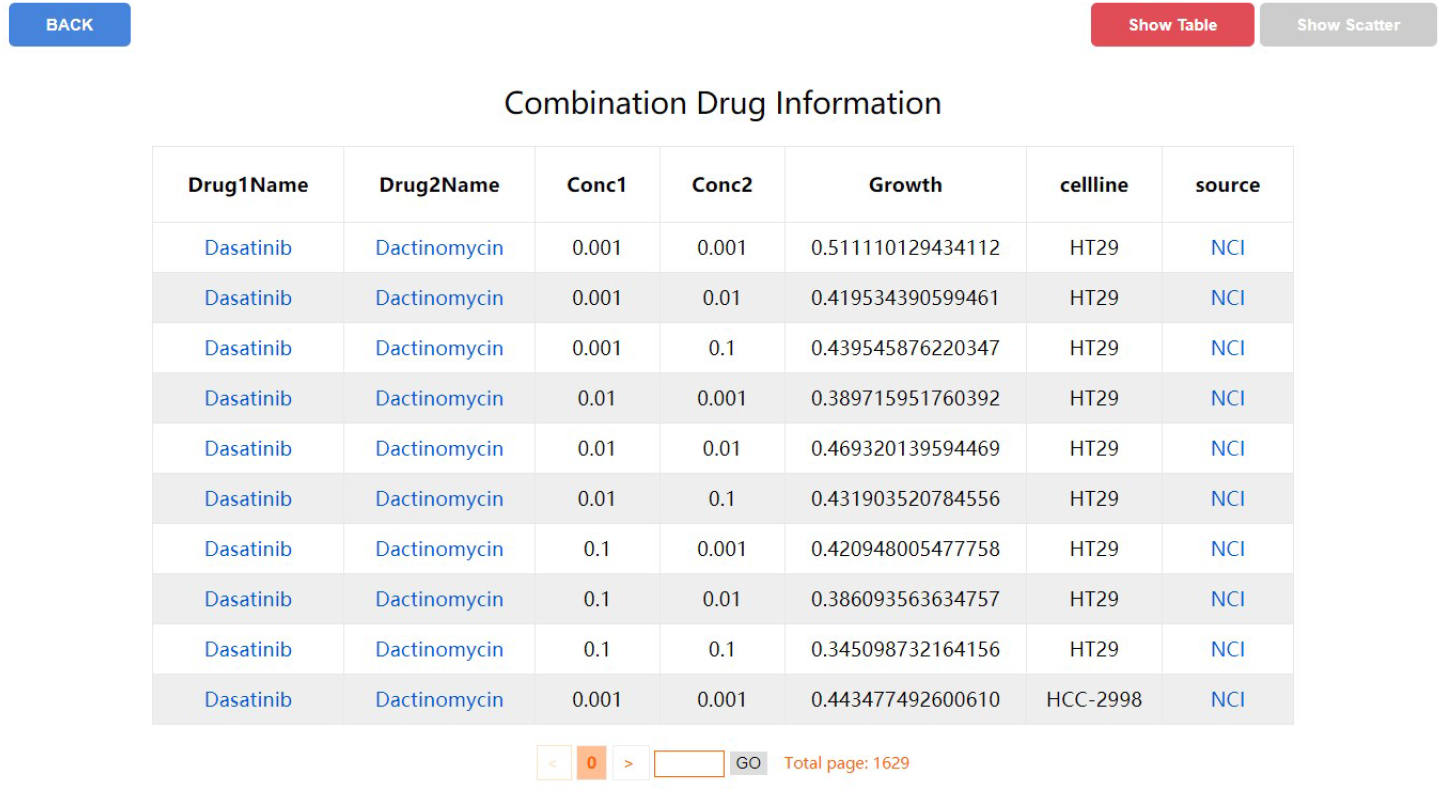
The predicted sensitivities of the combination drugs Lapatinib and Rapamycin.

